# Divergent stem cell mechanisms governing the primary body axis and appendage regeneration in the axolotl

**DOI:** 10.1101/2025.02.11.637618

**Authors:** Liqun Wang, Li Song, Chao Yi, Jing Zhou, Zhouying Yong, Yan Hu, Xiangyu Pan, Na Qiao, Hao Cai, Wandong Zhao, Rui Zhang, Lieke Yang, Lei Liu, Guangdun Peng, Elly M Tanaka, Hanbo Li, Yanmei Liu, Ji-Feng Fei

## Abstract

Exploring the fundamental mechanisms of organ regeneration is crucial for advancing regenerative medicine. The axolotl tail represents a unique opportunity to study regeneration of the primary axis including segmented muscle, vertebrae and skin. During tail development, muscle stem cells (MuSCs) displayed expected specificity to the muscle lineage. Tail amputation, however, induced expansion of MuSC potential yielding clonal contribution to muscle, connective tissue including cartilage, pericytes, and fibroblasts. This expanded potential was not observed during limb regeneration, and cross-transplantation showed these differences in potential are intrinsic. ScRNA-Seq profiling revealed that tail MuSCs revert to an embryonic mesoderm-like state. Through genetic manipulation involving the over-expression of constitutively active TGF-β receptors or Smad7 (antagonist of TGF-β signaling) in MuSCs, we demonstrated that the levels of TGF-β signal determine the fate outcome of MuSCs to connective tissue lineage or muscle respectively. Our findings illustrate a fundamental difference between regeneration of primary axis versus limb and offers a novel stem cell source for regeneration of axial skeletal tissues.

## INTRODUCTION

The axolotl regenerates multiple, complex organs, including the tail and limb^1–4^. These consist of similar mesodermal tissues including muscle, dermis, and skeleton but it is unknown whether the same stem cell mechanisms are implemented to regenerate these two body regions. Previous studies in the limb found that lineage-restricted progenitor/stem cells contribute to the blastema and give rise to corresponding tissues in regeneration^5,6^. The tail is of particular interest as it represents the regeneration of the primary axis. Amputation of the tail results in the regeneration of a functional structure that consists of all primary body axis tissues, including muscle, connective tissues (CTs) and spinal cord^7,8^. However, in comparison to the limb, our understanding of the cell origins of tail regeneration remains limited.

Muscle stem cells (MuSCs) are generally considered a unipotent stem cell that regenerates muscles after injuries, although a contribution to other cell types is sometimes observed^9–13^. In axolotls, fate mapping of genetically labeled *Pax7*+ MuSCs or labeled muscle tissue (muscle plus MuSCs) has consistently demonstrated their contribution exclusively to muscle during limb regeneration^6,14,15^. The tail myotomes also harbor *Pax*7+ MuSCs^8,14,16^ but their fate during tail regeneration has not been studied. From a developmental perspective, limb and tail MuSCs originate separately from the ventrolateral and dorsomedial portions of early somites^17–20^. They then migrate into the lateral plate or are situated in somites, giving rise to abaxial (limb) and primaxial (tail) muscles, respectively^19^. The primary body axis is considered an ancient feature that evolutionarily emerged much earlier than the appendage/limb^21,22^ and its regeneration has been observed in invertebrate chordate amphioxus^23^. Here we followed the fate of *Pax7*+ MuSCs, including clonal analysis during axolotl tail regeneration and surprisingly found that the tail MuSCs, in contrast to their limb counterparts, expand their potency to contribute to multiple CT lineages via dedifferentiation to an embryonic-like mesodermal progenitor state. The levels of TGF-β signaling regulate the fate outcome of MuSCs during tail regeneration, linking an injury signal with reversion to an early multipotent mesodermal cell-like state.

## RESULTS

### *Pax7+* cells produce connective tissue lineages during tail regeneration but not during development in juvenile axolotls

To identify the cell source contributing to the primary body axis during tail regeneration, we first bred *Pax7:CreER*^*T*2^ transgenic axolotls, in which expression of an inducible *Cre* is driven by the endogenous *Pax7* promoter^14^, with *CAGGS:reporter* line^24^ to establish *Pax7:CreER*^*T*2^*/CAGGS:reporter* double transgenic (dTG-*Pax7*) line (**Figure 1A**). Removal of the floxed-stop cassette upon tamoxifen administration in 2.5 cm dTG-*Pax7* axolotls, leads to *Cherry* expression allowing labeling and tracing of *Pax7+* cells and their progeny. After 14 days or 6.5 months tracing, the tamoxifen-converted CHERRY signal was restricted to the dorsal spinal cord and muscle compartments of dTG-*Pax7* axolotls as expected (**Figures 1B, 1C, 1F, 1G, S1A, and S1C**). Immunohistochemistry on sections showed that, apart from the CHERRY-converted spinal cord cells, the rest converted cells were positive for markers PAX7, MEF2C and MHC, revealing their muscle lineage identity^15^ (**Figure S2**).

**Figure 1.**
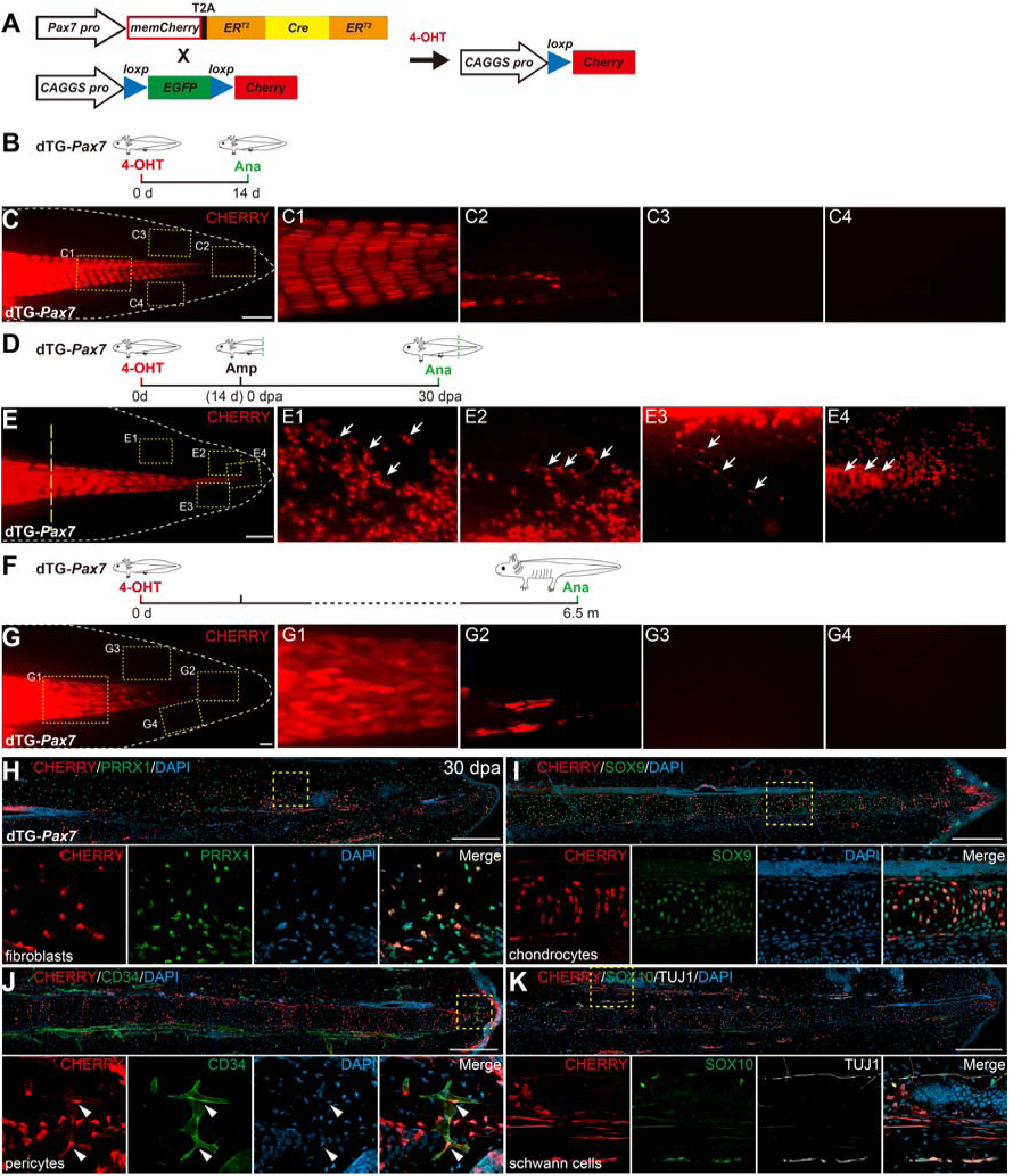
*Pax7+* cells give rise to multi-lineages during juvenile axolotl tail regeneration. (A) Strategy for genetic tracing of *Pax7+* cells in *Pax7:CreER^T2^/CAGGS:reporter* double transgenic (dTG-*Pax7*) axolotls. (B, D, F) Scheme of lineage tracing of *Pax7+* cells of 2.5 cm dTG-*Pax7* axolotls during development (B, F) and regeneration (D). (C, E, G) *Pax7*+ cells produce multi-lineages during tail regeneration, but not during tail growth. Representative images of CHERRY fluorescence in the tail of dTG-*Pax7* axolotls at 14 days (C, n=30), 6.5 months (G, n=6) after tamoxifen treatment, or 30 days post-amputation (dpa) following tamoxifen treatment (E, n=30). (H-K) Molecular characterization of the progeny of *Pax7*+ cells. Immunofluorescence for CHERRY (red) and PRRX1 (green, in H), SOX9 (green, in I), CD34 (green, in J), SOX10 and TUJ1 (green and white, in K), combined with DAPI (blue) staining on longitudinal sections of 30-day tail regenerates from the tamoxifen-converted dTG-*Pax7* axolotls (n=6). The rectangles (in C, E, G-K) are depicted at a higher magnification in their respective primed counterparts. Yellow dashed lines, the plane of amputation; white dashed lines, shape of the tail. Arrows, CHERRY+ progeny from converted *Pax7*+ cells in tail regenerates; arrowheads, CHERRY+ pericytes. Scale bars, 2 mm in C, G; 1 mm in E; 500 μm in H-K.

To map the fate of *Pax7*+ cells during regeneration, we amputated the tails of tamoxifen-converted dTG-*Pax7* axolotls and followed the progeny of CHERRY+ cells (**Figure 1D**). We observed that, in addition to the spinal cord and muscle, abundant CHERRY+ cells appeared throughout the regenerated fin and cartilage tissues at 30 days post-amputation (dpa) (**Figures 1E and S1B**). These CHERRY+ cells exhibited a typical morphology of fibroblasts, pericytes, chondrocytes of cartilage, and Schwann cells wrapping around peripheral nerves (**Figures 1E1-1E4 and S1D-S1G**). We further characterized the markers associated with converted CHERRY+ cells on longitudinal sections of 30-day tail regenerates by immunohistochemistry, and confirmed that converted *Pax7*+ cells gave rise to CHERRY+ peripheral nerves, Schwann cells, fin fibroblasts, chondrocytes, and pericytes, identified by the expression of markers TUJ1, SOX10, PRRX1 and SOX9, or by their physical locations (**Figures 1H-1K)**^24–27^. Examination of limb regenerates of dTG-*Pax7* axolotls by immunohistochemistry confirmed previous observations^6,14^ that converted CHERRY+ cells exclusively contributed to muscle, but not CT lineages during limb regeneration (**Figure S3**). These data show that the acquisition of extra CT lineages by *Pax7*+ cells is exclusive to tail regeneration, with no instances observed during tail and limb development or limb regeneration in juvenile axolotls.

### Muscle stem cells are the source contributing to connective tissues during tail regeneration

During development, *Pax7* expression is associated not only with myogenic lineages but also with the dorsal neural tube and neural crest^14,28–32^. We next sought to pinpoint the *Pax7* cell type that generates multi-lineages during tail regeneration. Without amputation, CHERRY-converted cells in dTG-*Pax7* axolotls include neural stem cells (NSCs), MuSCs, and their derived muscles (**Figures S2A and S2B**). We therefore created various *CreER^T2^* transgenic lines to trace these relevant cell types. Fate mapping of NSCs in the nervous system or mature muscles, using newly established *Sox2:CreER^T2^*/*CAGGS:reporter* or *MCK:CreER^T2^*/*CAGGS:reporter* double transgenic lines, showed that neither *Sox2*+ NSCs nor *MCK*+ muscle fibers gave rise to CT or muscle lineages during tail regeneration (**Figures S4 and S5**). We found Schwann cells contribution from CHERRY-converted *Sox2*+ NSCs (**Figure S4G**). These results suggest that the previously observed Schwann cells are likely derived from *Pax7*+ NSCs, while CT lineages potentially originated from *Pax7*+ MuSCs in dTG-*Pax7* axolotls.

To precisely determine the origin of CHERRY+ CT cells in dTG-*Pax7* axolotls during tail regeneration, we chose *Myf5*, another marker expressed in myogenic stem/progenitor cells^33–36^, and generated *Myf5:CreER^T2^*transgenic axolotls. Tracing the progeny of *Myf5*+ cells in 2.5 cm *Myf5:CreER^T2^*/*CAGGS:reporter* double transgenic axolotls (dTG-*Myf5*) revealed that converted CHERRY+ cells produced *Pax7*+ MuSCs and MHC+ muscles, but not spinal cord NSCs, 14 days after tamoxifen treatment in the uninjured tail (**Figures 2A, 2B, and S6**). However, following amputation, these CHERRY+ cells gave rise to CT lineages, including pericytes, PRRX1+ fibroblasts and cartilage chondrocytes in 30-day tail regenerates of dTG-*Myf5* axolotls (**Figures 2B-2D2**). These results demonstrate that *Myf5*+ myogenic stem/progenitor cells are capable of being committed to multi-lineages, and together with the *Pax7* tracing data, implying that MuSCs are the potential source giving rise to CT lineages during tail regeneration.

**Figure 2.**
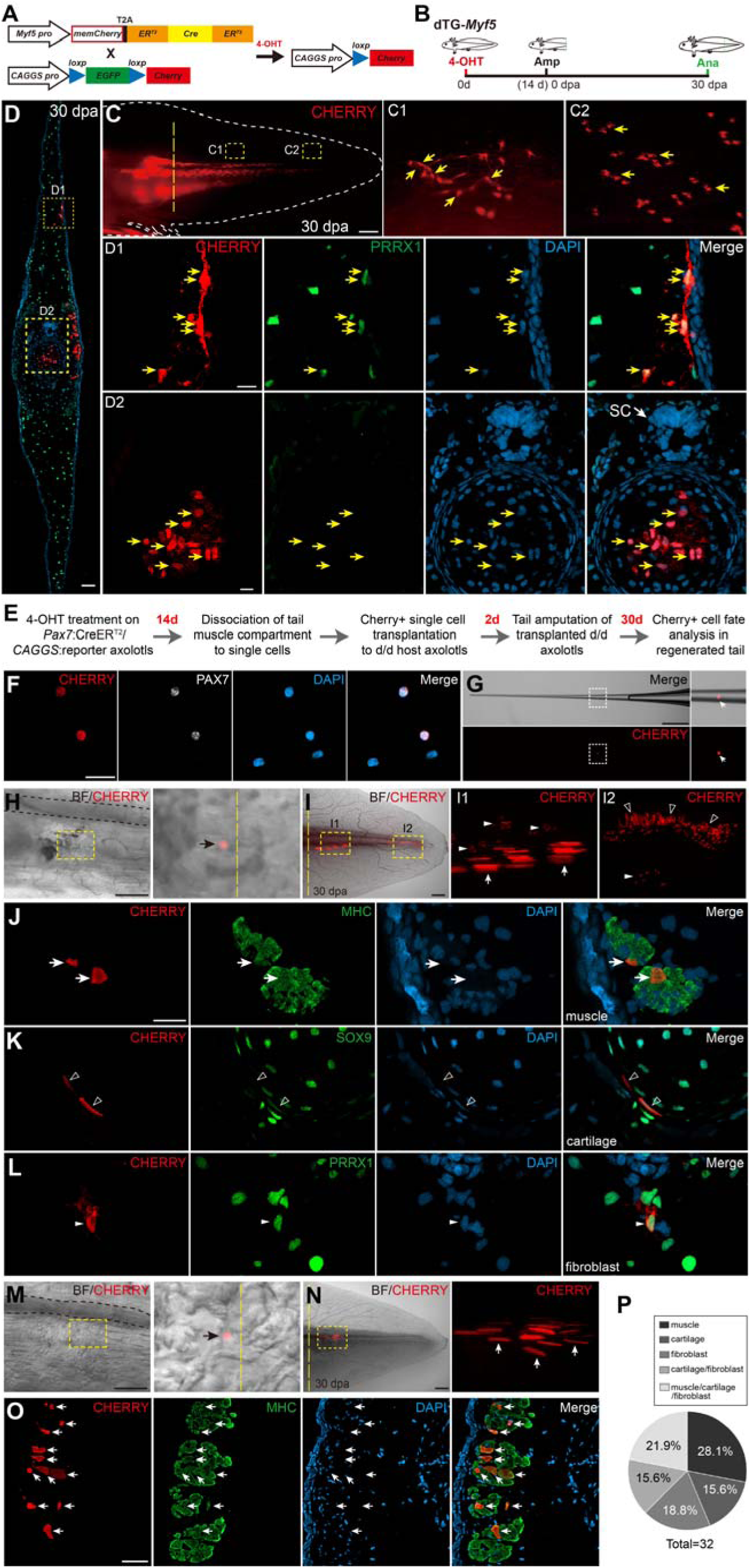
Muscle stem cells give rise to connective tissue lineages during tail regeneration. (A and B) Strategy (A) and experimental design (B) for genetic labeling and tracing of *Myf5+* cells in *Myf5:CreER^T2^*(F0)/*CAGGS:reporter* double transgenic (dTG-*Myf5*) axolotls. (C and D) Tamoxifen-converted *Myf5*+ cells give rise to connective tissues during tail regeneration. Representative images of CHERRY fluorescence in the tail (C) and immunofluorescence for CHERRY (red), PRRX1 (green), combined with DAPI staining on tail cross-sections (D) of tamoxifen-converted dTG-*Myf5* axolotls at 30 days post-amputation (dpa) (n=2). SC, spinal cord in D2. (E) Scheme of single muscle stem cell transplantation and fate mapping during tail regeneration. (F) PAX7 expression in isolated CHERRY+ single cells. Immunofluorescence for CHERRY (red) and PAX7 (white), combined with DAPI staining on isolated single cells from muscle compartments of tamoxifen-converted dTG-*Pax7*. (G) Merged (upper panels) and CHERRY-only (lower panels) fluorescence images of a single CHERRY+ cell loaded into a capillary pipette for transplantation. (H-O) Transplantation, and fate mapping of isolated single CHERRY+ cells during tail regeneration. Bright field (BF), CHERRY fluorescence, and merged live images showing a single CHERRY+ cell settling in the tail tissue of host *d/d* axolotls post-transplantation (H, M) and its derived progeny in the 30-day regenerated tail (I, N). J-L and O, Molecular characterization of the progeny of transplanted single cells shown in I and N. Immunofluorescence for CHERRY (red) and MHC (green, J and O), SOX9 (green, K), PRRX1 (green, L), combined with DAPI (blue) staining on cross-sections of 30-day regenerated tails from single cell-transplanted *d/d* hosts (n=32). (P) Quantification of transplanted single muscle stem cell fates during tail regeneration. The rectangles are depicted at a higher magnification in their respective primed counterparts. Yellow, white, and black dashed lines indicate the plane of amputation, shape of the tail, and spinal cord, respectively. Black arrows, transplanted single cells; white arrows, CHERRY+ muscles; white arrowheads, CHERRY+ fibroblasts; empty arrowheads, CHERRY+ chondrocytes. Scale bars, 2 mm in C; 1mm in G, I, N; 500 μm in H and M; 100 μm in D; 50 μm in F, J-L and O; 25 μm in D1-D2.

It has been reported that MuSCs are a heterogeneous population^9,37–39^. Live imaging revealed that tamoxifen-converted CHERRY+ cells in dTG-*Pax7* axolotls actively migrated into the newly formed tail blastema during regeneration, giving rise to distinct clusters potentially resembling clones of muscles and CTs (**Figure S7**). To investigate the potency and heterogeneity of *Pax7*+ MuSCs, we conducted single-cell transplantation^12^ and mapped their lineage during tail regeneration (**Figure 2E**). We dissociated tail muscle tissue from tamoxifen-converted dTG-*Pax7* axolotls to isolate CHERRY+ single cells, of which 87.7% (93/106) were positive for PAX7 by immunostaining (**Figure 2F**). A freshly isolated single CHERRY+ cell was then transplanted to the tail of a host *d/d* axolotl to trace its lineage during regeneration (**Figures 2G, 2H, 2M, S8A, S8D, and S8G)**. Following amputation, 16% (32/200) of transplanted host axolotls contained CHERRY+ cells in 30-day tail regenerates. Live imaging and immunohistochemical analysis on cross-sections of the tail regenerates revealed contributions to single or multiple lineages of muscle, chondrocytes, and fibroblasts (**Figures 2I-2L, 2N-2O, S8B-C, S8E-F, and S8H-J**). Out of the 32 successfully transplanted hosts, 21.9% (7/32) contributed to multiple lineages of muscle, cartilage, and fibroblast; 15.6% (5/32) contributed to cartilage and fibroblast; while 28.1% (9/32), 15.6% (5/32) and 18.8% (6/32) only muscle, cartilage, or fibroblast respectively (**Figure 2P**). These results demonstrate that tail *Pax7*+ MuSCs or their derived progenitors involved in regeneration are heterogeneous, with various differentiation potencies, showing myogenic or CT-genic unipotency, or dual myogenic and CT-genic multipotency.

Considering the differences observed in MuSCs between the tail and limb during regeneration, we next investigated the lineage and potency of limb MuSCs at the single-cell level. To this end, we carried out single-cell grafting of freshly isolated limb MuSCs to the tail, or tail MuSCs to the limb of d/d hosts and followed the fate of the transplanted cells (**Figure 3A**). We found that all grafted limb MuSCs (11/11) exclusively produced muscles, even in the tail regeneration environment (**Figure 3B**). These data further revealed that MuSCs in the limb, unlike their tail counterparts, are unipotent. Interestingly, we found that tail MuSCs gave rise to both muscle and CTs in the limb environment during regeneration (**Figures 3C and 3D**). These experiments indicate that the variations in differentiation potency in MuSCs are intrinsically determined, and MuSCs keep a memory of their original differentiation potency during regeneration, even when situated in a different environment.

**Figure 3.**
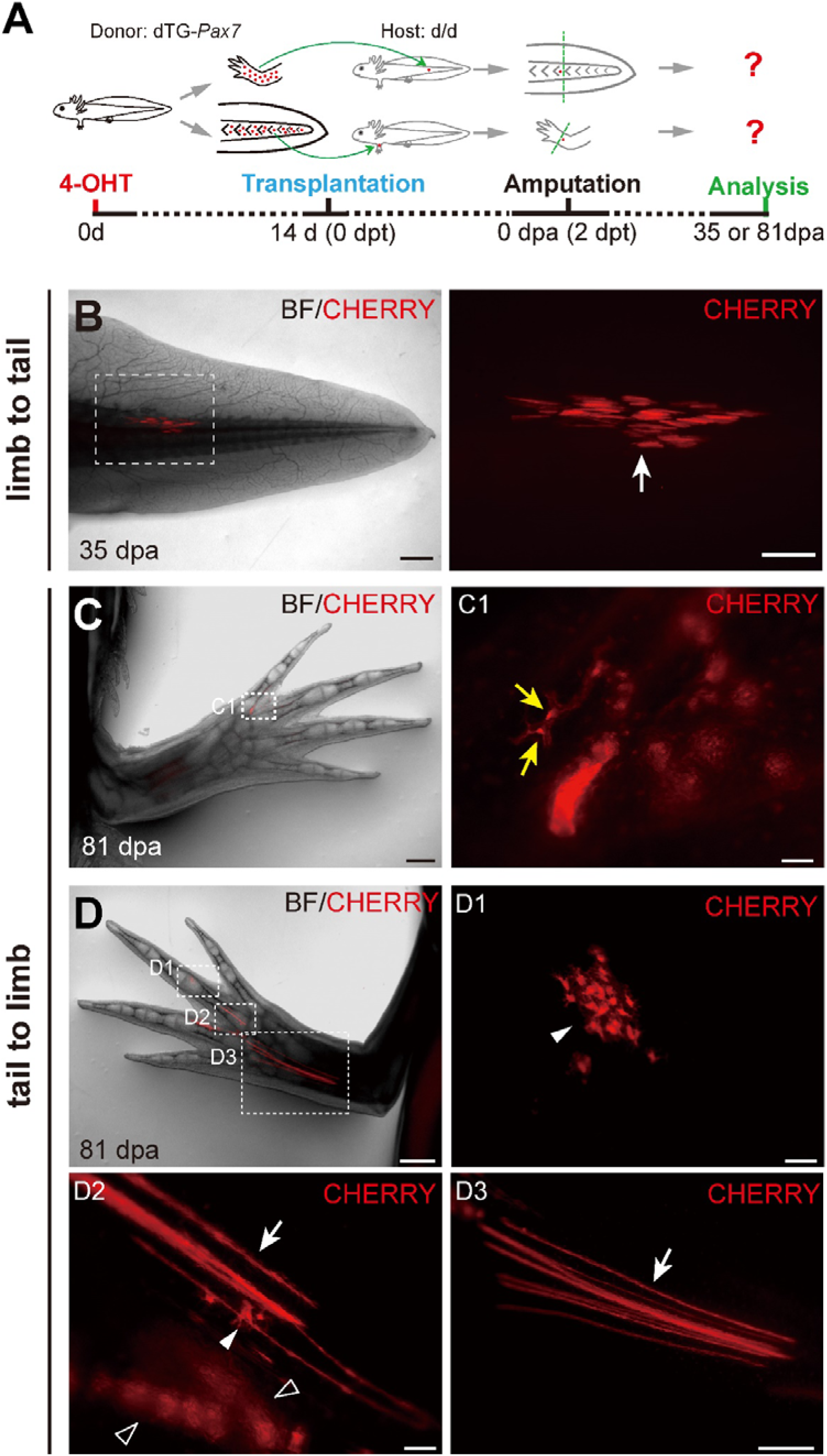
Reciprocal grafting of limb and tail MuSCs reveals potency variations. (A) Scheme for reciprocal grafting of single converted CHERRY+ satellite cells between the limb and tail, and fate mapping during limb and tail regeneration. Host *d/d* axolotls were amputated 2 days post-transplantation (2 dpt) of tamoxifen-converted, single CHERRY+ satellite cell, and analyzed at 35 or 81 days post-amputation (dpa). (B-D) Bright field (BF) and CHERRY fluorescence images showing the progeny of reciprocal transplanted MuSCs (limb to tail, B; tail to limb, C and D) during regeneration. Note that a single grafted limb MuSC produces only muscle lineages (n=11) during tail regeneration (B). However, a single grafted tail MuSC gives rise to both muscle and connective tissue lineages (fibroblasts and chondrocytes, n=3) (D), or only connective tissue lineages (pericytes and chondrocytes, n=7) (C) during limb regeneration. The rectangles are shown at higher magnification as single-channel CHERRY images. Solid and empty arrowheads indicate fibroblasts and chondrocytes; white and yellow arrows indicate muscle fibers and pericytes, respectively. Scale bars, 2 mm in b; 1 mm in C, D, B1; 500 μm in D3; 100 μm in C1, D1, D2.

### Muscle stem cells give rise to connective tissue lineage through intermediate progenitors

To elucidate the transition routes underlying the multipotency acquisition of *Pax7+* MuSCs during axolotl tail regeneration, we collected tamoxifen-converted CHERRY+ cells and their progeny by FACS from uninjured and regenerating tails of dTG-*Pax7* axolotls (**Figure S9A**), and carried out single-cell RNA Smart-seq2 sequencing (scRNA-seq)^40^. We obtained a total of 1640 cells, with an average of 7454 genes detected per cell (**Figure S9B**).

We then analyzed scRNA-seq data by Louvain clustering^41^ and annotated 20 subtypes, including cell types in the muscle and neuronal lineages such as MuSCs (*Pax7*, *Myf5*), myoblasts (*Myog*, *Myf5*), neural progenitors (*Pax7*, *Gfap*, *Sox2*), Schwann cells (*Sox10*) and neurons (*Neurod6*); cell types in the CT lineages such as fibroblasts (*Prrx1*, *Pdgfra* and *Lum*) and chondrocytes (*Sox9*, *Pdgfra* and *Lum*) (**Figures 4A, 4B, and S9C**). Remarkably, we also uncovered two subtypes that exhibited great similarity to fibroblasts (*Prrx1*, *Pdgfra* and *Lum*) but also expressed MuSC markers *Pax7* and *Myf5* (**Figures 4A and 4B**), therefore named as intermediate cell cluster 1 (IM1) and 2 (IM2). We separated cells from all samples into three compartments on UMAP according to gene expression similarity (**Figure S9D**), named “region 1-3” (**Figure 4C**). MuSCs, IMs, and fibroblasts fell into “region 1”. To explore the temporal dynamics of the identified cells throughout tail regeneration, we further subdivided seven samples into four phases according to the kinetics of MuSCs and IMs (**Figure 4C**). We found that prior to amputation, consistent with previous fate mapping data, nothing other than muscle lineage existed in “region 1” at phase I (0 dpa) (**Figure 4C**). IM1 and IM2 sequentially emerged at phase II (2, 4, 6 dpa) and III (10, 15 dpa), concomitant with a decline of MuSCs and myoblasts, and a burst of fibroblasts and chondrocytes, accordingly (**Figure 4C**). At Phase □ (33 dpa), close to the completion of tail regeneration, MuSCs and myoblasts in “region 1” reverted to a state resembling that at 0 dpa, coinciding with the vanishing of IMs (**Figure 4C**). We therefore designated “region 1” as the “regeneration active zone”.

**Figure 4.**
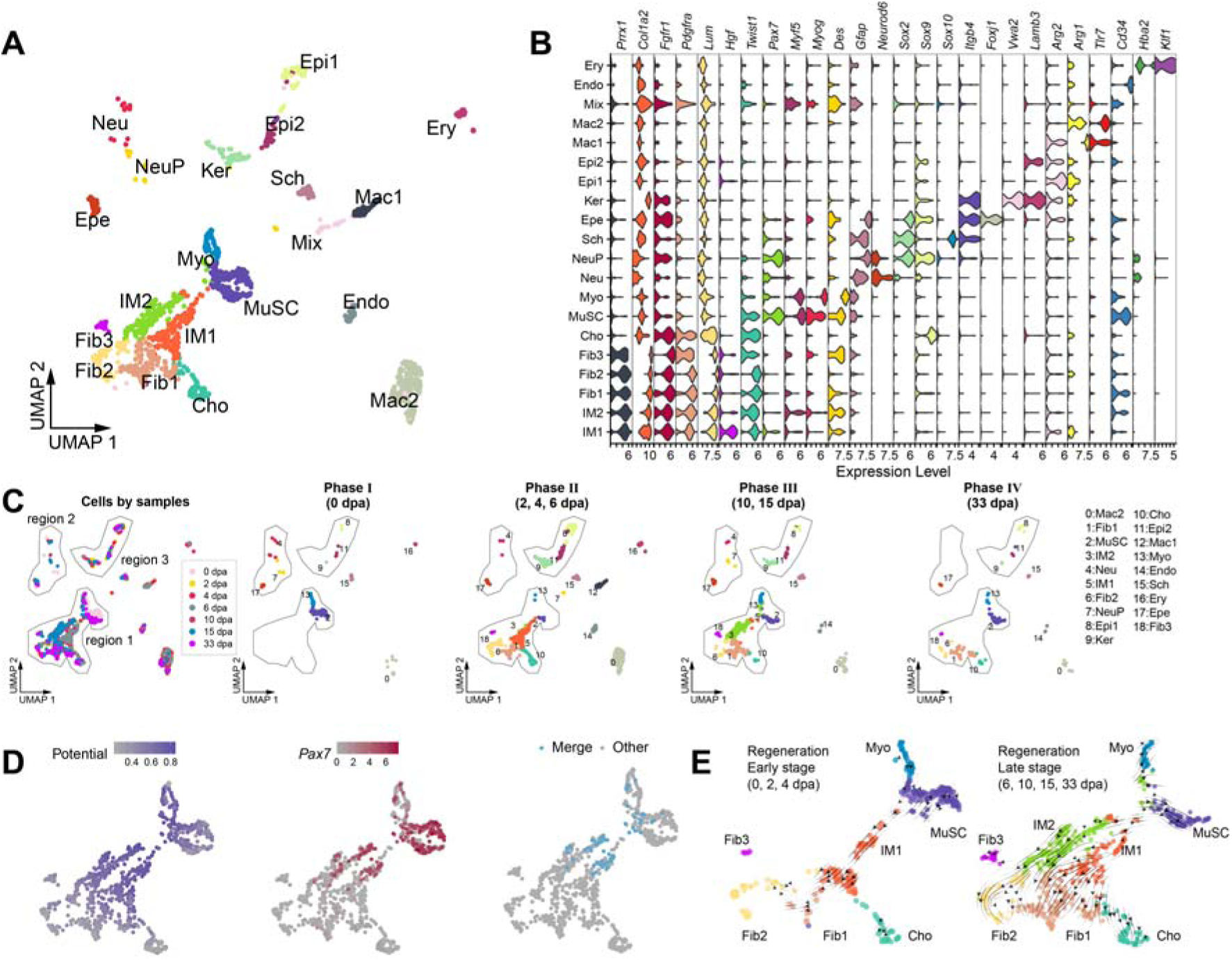
Muscle stem cells give rise to connective tissue lineages via intermediate progenitors during tail regeneration. (A) Uniform manifold approximation and projection (UMAP) of single-cell clustering from Smart-seq2 data. 20 cell types were annotated, including Cho, Chondrocyte; Sch, Schwann cell; Endo, Endothelial cell; Ery, Erythroid Cell; Epe, Ependymoglial cell; Epi1, Epithelial cell 1; Epi2, Epithelial cell 2; Fib1, Fibroblast 1; Fib2, Fibroblast 2; Fib3, Fibroblast 3; IM1, Intermediate cell 1; IM2, Intermediate cell 2; Ker, Keratinocyte; Mac1, Macrophage 1; Mac2, Macrophage 2; MuSC, Muscle stem cell; Myo, Myoblast; Neu, Neuron; NeuP, Neural progenitor cell, with one unknown cell type designated as “Mix”. (B) Violin plot displaying expression levels of representative markers in each annotated cell type. (C) UMAP visualization of cells from all sampling stages (left) and dynamics of cell types at different Phases (right). Three regions with lineage-related cells are delimited by polygons and labeled as “region 1-3”. Note that major activities occur in “region 1”, designated as the “regeneration active zone”. (D) Heatmaps displaying stemness potential (left), *Pax7* expression levels (middle), and the merge of top 20% stemness potential and *Pax7* expression levels >= 3 (right) in cells within the “regeneration active zone”. (E) RNA velocity streamline plots predicting lineage transitions among myogenic-related and connective tissue cells at early (left, 0, 2, 4 dpa) and late (right, 6, 10, 15, 33 dpa) regeneration stages, respectively.

To determine the origin and descendants of IMs in regeneration, we performed a stemness score calculation^42^ and uncovered that IM1 and IM2, similar to MuSCs, exhibited a higher stemness score and *Pax7* expression (**Figures 4D and S9E**). We then performed RNA velocity analysis^43^ and revealed a transition from MuSCs to IM1 at the early stage of regeneration (**Figure 4E left**). Trajectory analysis^44^ also showed that MuSCs differentiated into myoblasts before amputation (0 dpa) but switched to IM1 during early regeneration (2, 4 dpa) (**Figure S10**). The emergence of IM1 appeared to occur at the expense of MuSCs, as evidenced by a dramatic decline of MuSCs and myoblasts (**Figure 4C**). Whereas, at the late stage of regeneration, we noticed a shift from IM1 to IM2, followed by differentiation into CT cells or replenishment of the consumed MuSCs (**Figures 4C and 4E right**).

In conclusion, our data indicate that IMs, as a regeneration-induced stem/progenitor population, are derived from MuSCs and act as the direct source contributing to CT lineages during tail regeneration.

### The state and lineages of IM progenitors resemble that of early embryonic multipotent mesodermal cells

We next assessed the ground state of MuSC-derived IMs. Previous reports have shown that limb fibroblasts, and brain ependymoglial cells in axolotls revert to a more primitive, embryonic-like state during regeneration^25,45,46^. Therefore, we wondered whether IMs share similar gene expression profiles and lineages with certain types of early embryonic cells, e.g., mesodermal cells. It has been reported that a subpopulation of *Pax7*-expressing mesodermal cells gives rise to trunk adipocytes, dermis, and muscles during early mouse embryogenesis (e.g., E9.5 and E10.5), but become lineage-restricted to generate only muscles as development proceeds (e.g., E12.5)^29,30^. Whole-mount in situ hybridization on early axolotl embryos and immunohistochemical analysis on sections showed *Pax7* expression in neural crest, neural tube, somitic and pre-somitic mesoderm cells (**Figures 5A and S11A**), consistent with observations in rodents^29,30^. We then treated dTG-*Pax7* embryos with a consecutive or single dose of tamoxifen at stages ranging from 15 to 44, to map the fate of *Pax7+* cells in the tail of 2 cm axolotls during development (**Figures 5B and S11B**). In addition to muscle and neuronal lineages, *Pax7*+ cells made significant contributions to CT lineages. CHERRY+ cells converted at stage 15 to 25 by a single pulse tamoxifen produced abundant CHERRY+ progeny, including dermis, fibroblasts, and pericytes in the fin tissue of axolotl larvae, while those converted at stage 30 or 44 yielded fewer or no CHERRY+ progeny (**Figures 5C and S11C-S11F**). This transient CT lineage commitment in *Pax*7+ cells during early embryogenesis might resemble the induction of CT lineages observed in MuSCs during juvenile axolotl tail regeneration. Collectively, our results suggest that a population of *Pax7+* embryonic progenitor cells (EPrCs) within the neural crest or mesoderm are able to transiently give rise to CT lineages during axolotl embryonic tail development.

**Figure 5.**
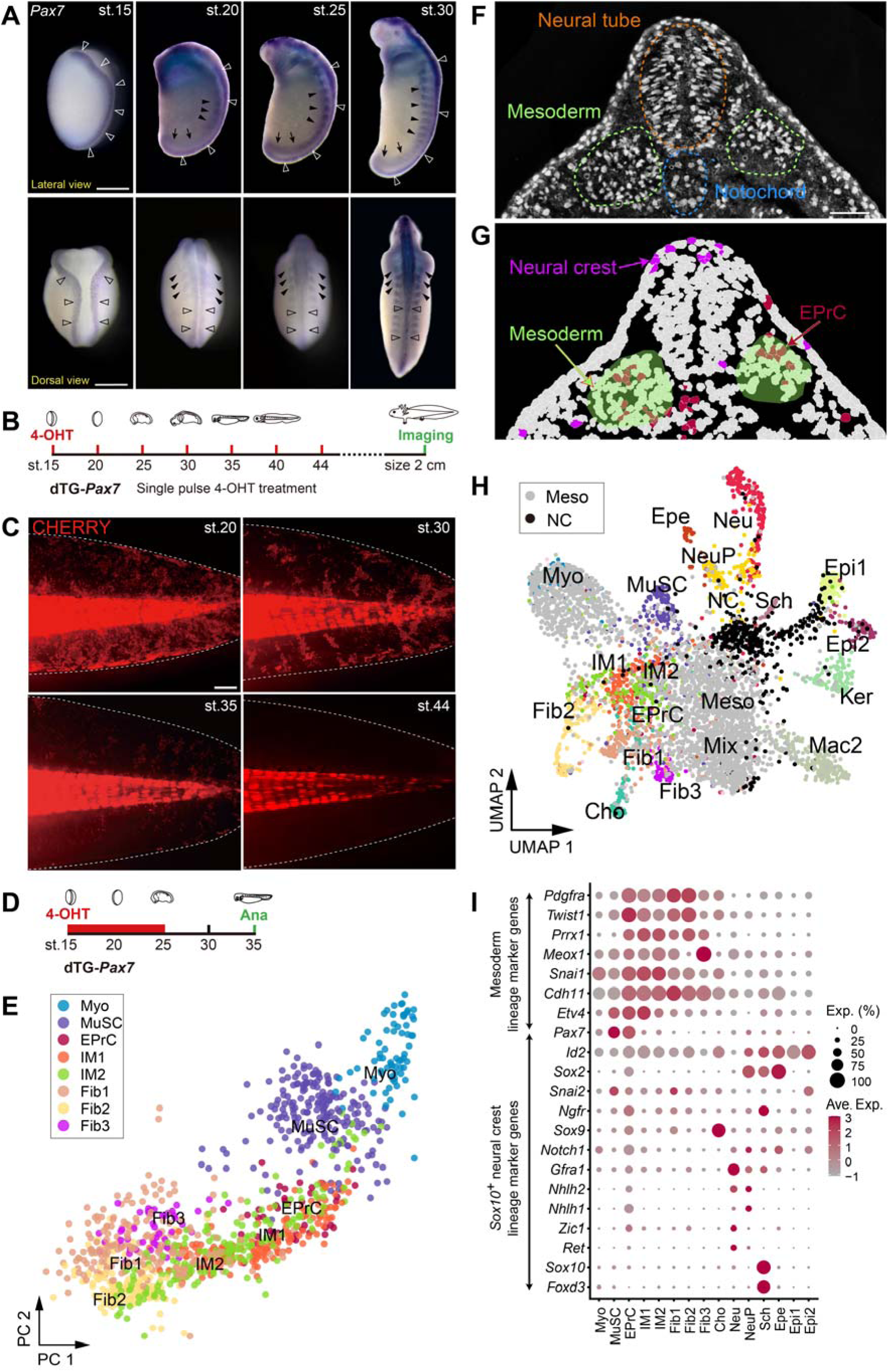
The state and lineages of intermediate cells (IMs) resemble that of early embryonic multipotent mesodermal cells. (A) Embryonic expression of *Pax7*. Whole-mount in situ hybridization of axolotl embryos at stage (st.) 15, 20, 25, 30. Empty arrowheads, arrowheads, and arrows indicate *Pax7* expression in the neural crest, somite, and pre-somatic mesoderm, respectively. (B) Scheme for genetic tracing of *Pax7+* cells at early embryonic stages. dTG-*Pax7* embryos were treated with a single dose of tamoxifen at the indicated stages, and analyzed in 2 cm axolotls. (C) Lineages of early embryonic *Pax7*+ cells. Representative CHERRY fluorescence images of tails from 2 cm dTG-*Pax7* axolotls treated with tamoxifen at indicated stages. (n=3-5 embryos per stage). (D) Scheme of Smart-seq2 sample collection at embryonic development stages. (E) PCA plot showing cell type similarity. EPrCs overlay with IMs, which indicates IMs from regeneration resembling the state of EPrCs from embryos. EPrC, early embryonic progenitor cell. (F and G) Spatial visualization of EPrCs on Stereo-seq section of stage 25 axolotl embryos. Dark red, EPrC scRNA data projected to Stereo-seq data; Magenta, *Sox10*+ neural crest cells; light green, mesoderm areas. Note that EPrC is mainly located in the mesoderm area as determined by ssDNA staining (F). (H) Integration of Smart-seq2 and embryonic Stereo-seq data. EPrCs and cells from the regeneration active zone primarily clustered with mesodermal cells, but not neural crest cells. Meso, mesoderm; NC, neural crest. (I) Bubble plot exhibiting the expression of neural crest and mesoderm marker genes in annotated cell types from Smart-seq2 data. Scale bar, 1 mm in A; 500 μm in C; 100 μm in F-G.

We next attempted to evaluate the likeness of EPrCs and IMs. To this end, we consecutively treated dTG-*Pax7* embryos with tamoxifen from stage 15 to 25, and isolated tamoxifen-converted CHERRY+ cells containing EPrCs at stage 35 for Smart-seq2 sequencing (**Figure 5D**). We then integrated and clustered this data with scRNA-seq data from regeneration samples, and uncovered three main cell types, including neural progenitors, neurons, and a new cell cluster from traced embryos. This new cluster was drawn in the “regeneration active zone”, and expressed *Pax7* and *Prrx1* (**Figures S12A-S12C**), speculating its EPrC identity. A constellation plot showed intensive interplays among EPrC, MuSC, and IMs, demonstrating shared expression profiles among these cell types (**Figure S12D**). Moreover, principal component analysis demonstrated an overlap between EPrCs and IMs on the plot (**Figure 5E**), suggesting that IMs partially recapitulate the state of EPrCs.

As previously observed, EPrCs that contributed to CT lineage were potentially derived from neural crest or mesodermal cells (**Figures 5A-5C and S11**). To define the origin and localization of EPrCs, we performed Stereo-seq on cross-section of stage 25 axolotl embryos and projected EPrCs to the spatial transcriptomics data^47^. Such projection showed that EPrCs were primarily located within the mesoderm region, as determined by ssDNA staining on the Stereo-seq section^48^ (**Figures 5F and 5G**). In addition, by extracting mesoderm and neural crest cells from the spatial transcriptomics data and utilizing integrative clustering on extracted spatial segmented cells and all scRNA data, we observed that IMs and EPrCs were predominantly situated in the embryonic mesoderm zone, separated from the *Sox10*+ neural crest area on the UMAP (**Figure 5H**). Consistently, marker genes relevant to the mesoderm^49^, but not neural crest cells^50^, were expressed in IMs and EPrCs (**Figure 5I**). Taken together, these data suggest that both IMs and EPrCs closely resemble the state of early mesodermal cells, implying that tail MuSCs revert to a multipotent embryonic mesodermal cell-like state to produce both muscle and CT cells during regeneration.

### TGF-β signaling pathway regulates the cell fate switch of myogenic MuSCs

To identify a molecular determinant modulating the cell fate switch of MuSCs towards CT lineages, we further sub-clustered MuSCs and IMs according to their differentiation potential (**Figure 6A**), and obtained nine subtypes committed to fibroblasts (IM1_Fib1, IM2_Fib2), muscle lineages (IMs_MuSC, IM2_Myo) or uncommitted. We then used these nine subtypes, combined with Fib1, Fib2 and Myo, major downstream cell types relevant to fibroblast or muscle lineages, to analyze pathway activity guiding the cell fate switch^51^. We observed that these cells were grouped into two segments (**Figure 6B**). IMs committed to fibroblast and muscle, situated in the upper and lower segments respectively, displayed the most divergent activity on the heatmap. Among them, the Wnt, TGF-β, and MAPK pathways emerged as the top three activated pathways.

**Figure 6.**
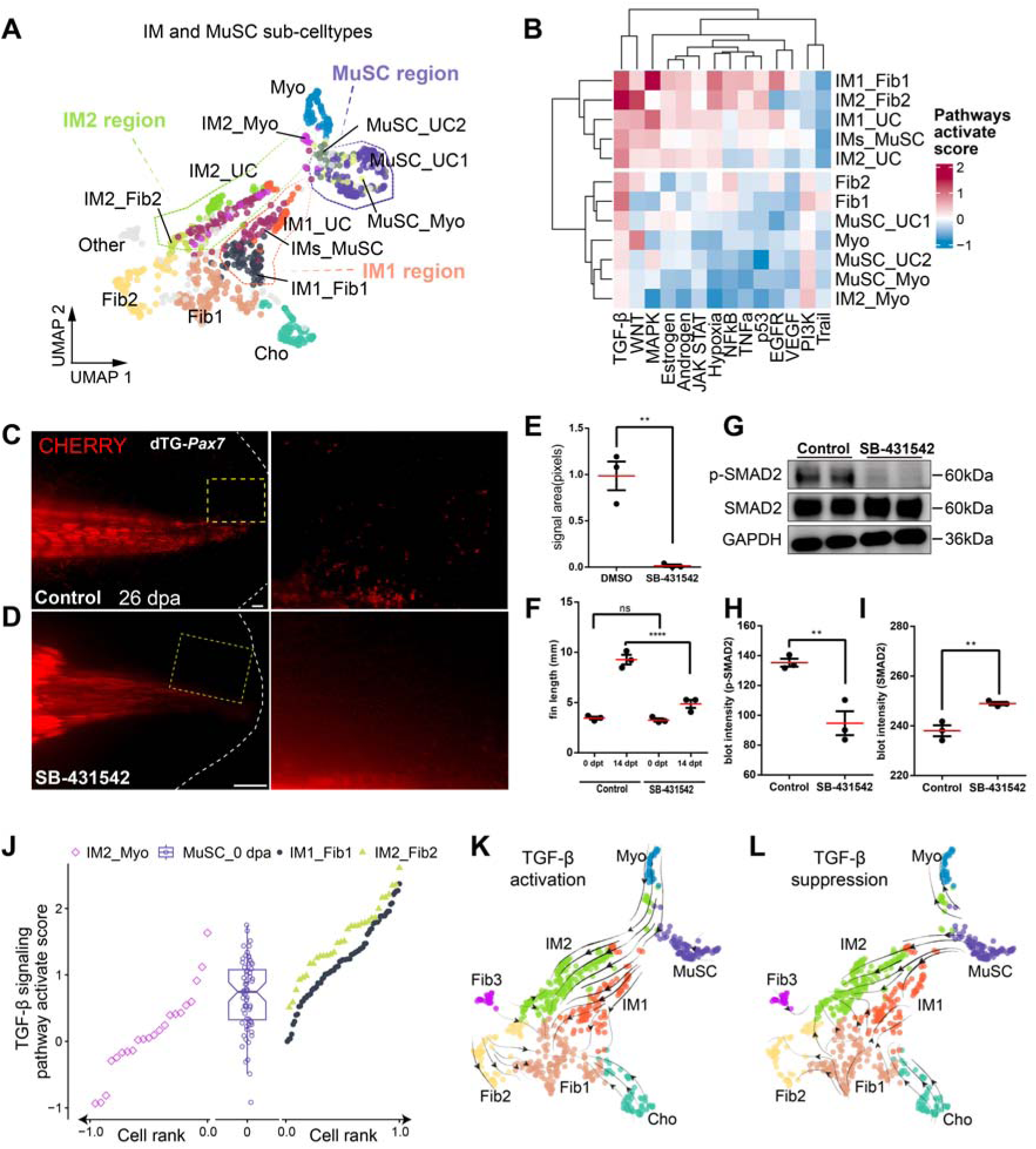
TGF-β signaling regulates the cell fate switch of muscle stem cells. (A) UMAP visualization of refined sub-clusters within the “regeneration active zone”. MuSC, IM1, and IM2 are partitioned into nine sub-clusters based on shared gene expression programs between progenitors and their progeny. Each cell type, e.g., IM2, is subdivided into sub-clusters committed to myoblasts (IM2_Myo), fibroblast 2 (IM2_Fib2), or uncommitted (IM2_UC). (B) Heatmap showing the scores of predicted pathway activity in nine defined sub-clusters from Figure 5a, combined with Fib1, Fib2, and Myo. (C and D) SB-431542 inhibits the production of fibroblasts during tail regeneration. Representative images of CHERRY fluorescence of 26-day tail regenerates treated with SB-431542 (D) and control (C) (n=3 each). (E and F) Quantification of CHERRY signal area on the fin and the length of the regenerated fin upon SB-431542 treatment (n=3). dpt, days post-treatment. Error bars, SEM; **p<0.01, ****p<0.0001; ns, not significant. (G) p-SMAD2 and SMAD2 immunoblots of regenerated tails treated with SB-431542 and control (n=3 each). GAPDH immunoblot, loading control. (H and I) Quantification of signal intensity from p-SMAD2 and SMAD2 immunoblots in (G) (n=3 each). Error bars, SEM; **p<0.01. (J) Boxplot of the scores of endogenous TGF-β activity in MuSCs. Note that scores are attenuated in IM2 committed to myoblasts (IM2_Myo), in contrast, elevated in IMs to fibroblasts (IM1_Fib1, IM2_Fib2), compared to that in MuSCs at 0 dpa. (K and L) Cell fate trajectory predictions following *in silico* perturbation of the TGF-β signaling pathway. Activation of TGF-β signaling diverts cells from muscle-related lineages to fibroblast-related lineages (K), whereas suppression of TGF-β signaling diverts cells from fibroblast-related lineages to muscle-related lineages (L). The rectangles (in C, D) are shown at higher magnification. Scale bars, 1 mm in C, D.

Based on the above analysis, we next experimentally investigated the roles of these pathways in regulating the cell fate switch of *Pax7*+ MuSCs during regeneration. We first applied drugs IWR-1^52^, SB-431542^53^, or U0126 ^54^ to dTG-*Pax7* axolotls to inhibit Wnt, TGF-β or MAPK pathways respectively, and traced the fates of tamoxifen-converted CHERRY+ cells during regeneration under each condition. We found that inhibition of the TGF-β pathway with SB-431542, evidenced by the down-regulation of phosphorylated SMAD2 (p-SMAD2) (**Figures 6G-6I**), significantly reduced the production of fibroblasts from CHERRY+ MuSCs and shortened the fin length during tail regeneration—a phenotype not observed upon the suppression of the Wnt or MAPK pathways (**Figures 6C-6F and S13**). In line with the drug inhibition results, bioinformatic analysis of TGF-β activity^43^ on the identified sub-clusters committed to different lineages revealed that the score of TGF-β signaling activation tended to increase in IM1_Fib1 and IM2_Fib2 committed to fibroblast lineage, but decrease in IM2_Myo committed to the muscle lineage, when compared to MuSC_0 dpa (**Figure 6J**). Furthermore, genetic perturbance analysis^43^ suggests that MuSC preferentially differentiates to fibroblast upon TGF-β activation (**Figure 6K**). In contrast, attenuation of TGF-β signaling promotes muscle lineage (**Figure 6L**). Collectively, our data suggest that TGF-β, rather than Wnt or MAPK, is a potential regulator of the MuSCs fate switch.

Considering the potential side effects of drugs on regeneration, we next carried out a genetic approach to manipulate TGF-β signaling. We expressed constitutively active (*ca*), dominant negative (*dn*) formats of *Tgfb receptor 1* (*Tgfbr1*)^55,56^ or *Smad7*, an antagonist of TGF-β signaling^57^, in *Pax7*+ cells to study their roles in tail regeneration. To this end, we first validated these constructs in cell culture. We found that transfection of plasmids encoding *Smad7*, but not that encoding *dnTgfbr1*inhibited TGF-β activity, while expression of *caTgfbr1* promoted TGF-β activity, indicated by the level of p-SMAD2 (**Figure S14**). We then created *Pax7:CreER^T2^*/*CAGGS:loxp-STOP-loxp-caTgfbr1/Smad7-Cherry* double transgenic axolotls (dTG-*caTgfbr1*; dTG-*Smad7*) (**Figure 7A)**, and investigated the cell fate commitment of *Pax7*+ MuSCs during tail regeneration, upon over-expression of *caTgfbr1* or *Smad7*. After tamoxifen treatment, we observed the CHERRY-conversion in the NSCs of the dorsal spinal cord and MuSCs (**Figures S15 and S16**), and the over-expression of *caTgfbr1* or *Smad7* in converted cells in dTG-*caTgfbr1* and dTG-*Smad7* lines, compared to the control dTG-*Pax7* axolotls (**Figure S17**). Following amputation, we noted that compared to the control (**Figures 7B and 7E**), when expressing *Smad7*, the production of CHERRY+ CT cells was nearly completely blocked, and only muscle lineage was visualized in the tail regenerates (**Figures 7C and 7F**). In contrast, the expression of *caTgfbr1* directed the converted CHERRY+ cells toward the CT rather than the muscle lineage (**Figures 7D and 7G**). These results demonstrate that TGF-β signaling alters the balance of the cell fate switch of MuSCs towards muscle or CT lineages during tail regeneration.

**Figure 7.**
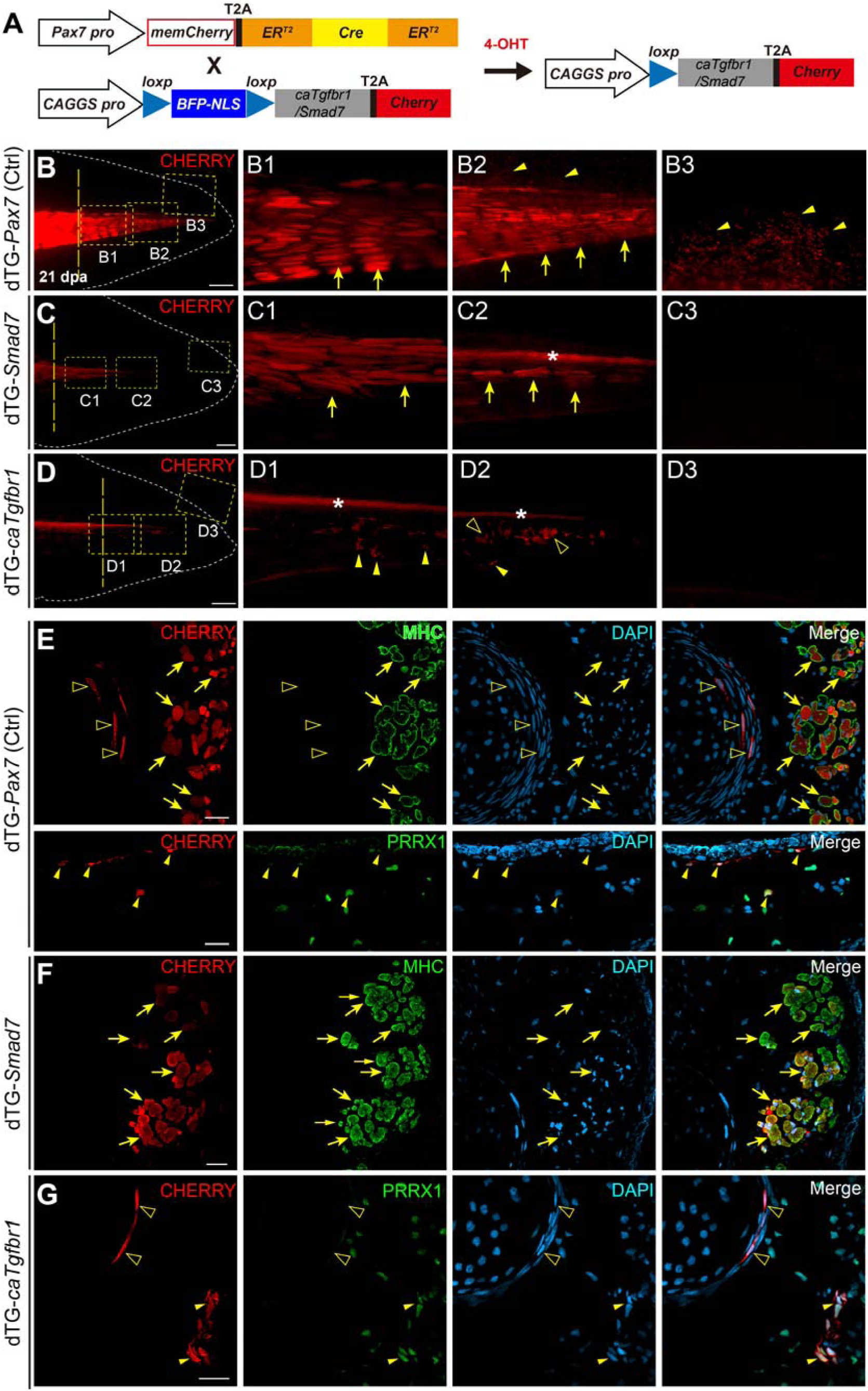
Modulating TGF-β activity leads to the cell fate switch of muscle stem cells during tail regeneration. (A) Strategy for modulating TGF-β activity in *Pax7*+ cells and cell fate mapping. In *Pax7:CreER^T2^*/*CAGGS:target-gene-reporter* (F0) double transgenic axolotls, named as dTG-*caTgfbr1* and dTG-*Smad7*, expression of constitutively active *Tgfb receptor 1* (*caTgfbr1*) or *Smad7*, along with the *Cherry* gene driven by the *CAGGS* promotor, are induced specifically in *Pax7+* cells after tamoxifen treatment. (B-D) Genetic inhibition or activation of *Tgfb* attenuates connective tissue or muscle lineages, respectively, in *Pax7*+ muscle stem cells during tail regeneration. Images of the progeny of tamoxifen-converted CHERRY fluorescence of 21-day tail regenerates upon expression of *caTgfbr1* (D), *Smad7* (C) and control (B). The rectangles are shown at higher magnification (in B1-D3). Note the complete loss of connective tissue or muscle lineages upon expression of *Smad7* or *caTgfbr1*, respectively. (E-G) Immunofluorescence for CHERRY (red), MHC (green), or PRRX1 (green), combined with DAPI staining, on cross-sections of 21-day tail regenerates of control (Ctrl) dTG-*Pax7*, dTG-*Smad7*, and dTG-*caTgfbr1* axolotls. Arrows, muscles; arrowheads, fibroblasts; empty arrowheads, chondrocytes; asterisks, spinal cord. Yellow and white dashed lines indicate the plane of amputation and the shape of the tails, respectively. B-D, n=3, 7, 10. Scale bars, 1 mm in B-D; 50 μm in E-G.

## DISCUSSION

Here we genetically traced the progeny of *Pax7*+ MuSCs, combined with fate mapping of transplanted single MuSC, and found differentiation potency varies in MuSCs during primary body axis and appendage regeneration. MuSCs in the axolotl tail revert to a multipotent state resembling embryonic mesodermal cells and give rise to muscle and CT lineages during regeneration. Furthermore, we showed that the multipotency commitment of MuSCs is modulated by TGF-β signaling levels, and demonstrated that controlling TGF-β activities led to the switch between muscle and CT lineages during tail regeneration.

What causes the difference in MuSCs between the tail and limb during regeneration? From an evo-devo perspective, the primary body axis (tail) and appendage (limb) exhibit the following variations. First, the establishment of the primary axis occurs as early as the gastrulation stage in most vertebrates^58,59^. In axolotls, the formation of the somites starts at about stage 20, followed by the appearance of a protrusion-like tail bud at stage 21. Whereas limb bud development is initiated at about stage 40^59^. Second, a muscle-containing appendage limb evolutionarily emerged much later than the primary body axis tail^21,22,60^. Consequently, the cells that contribute to tail and limb formation could be potentially different^18–20^. Indeed, MuSCs in the tail and limb originate from the dorsomedial and ventrolateral parts of the somites. Moreover, the muscle tissues, including MuSCs of the tail and limb, are embedded respectively in a presomitic mesoderm- and lateral plate mesoderm-derived CT environments^18–20^. These differences might be the primary rationale for our discovery that tail MuSCs, in contrast to their limb counterparts, contribute additionally to CT cells during regeneration. It will be important to further identify the key intrinsic or extrinsic factors resulting in the potency variations between tail and limb MuSCs during regeneration.

The multipotency of MuSCs was initially observed under in vitro cell culture conditions, where they were found to generate adipocytes and osteocytes, besides myocytes^61–64^. Subsequent studies revealed that MuSCs were also capable of producing similar multi-lineages during mouse embryonic development^13,29,30,65–68^, however, they seldom commit to multi-lineages spontaneously during homeostasis or regeneration^9^. In our study, by applying genetic fate mapping, particularly, tracing the progeny of freshly isolated single-transplanted MuSCs, which provides the most direct evidence of heterogeneity and multipotency, we observed a unique induced multipotency conversion of MuSCs during tail regeneration and a potency variation between limb and tail MuSCs. Interestingly, previous findings in newts showed that the transplantation of in vitro expanded MuSCs gives rise to adipocytes, chondrocytes and myocytes during limb regeneration^69^. In that study, MuSCs from newts were isolated in the absence of genetic labeling, and the experiment was not carried out at the single-cell level. Therefore, the purity of the MuSCs could not be precisely determined. Moreover, the MuSCs underwent in vitro expansion prior to transplantation, potentially altering their properties and complicating the interpretation of their in vivo behavior. Furthermore, despite both being salamanders, the axolotls and newts employ distinct strategies to regenerate limb muscles. Axolotls exclusively rely on resident MuSCs, whereas newts can regenerate muscle from progenitors derived from muscle dedifferentiation^15,70^. This divergence may endow the intrinsic differences in MuSCs between these two species. It is worth systematically investigating the differentiation potential of MuSCs between the limb and tail from the newt in development and regeneration environments.

Although early studies showed that elevated TGF-β signaling impairs myoblast differentiation^71,72^, little is known about its role in the cell fate commitment of MuSCs during regeneration. A recent study demonstrated that TGF-β signaling plays a negative role in muscle fusion during regeneration^73^. In our study, via analysis of scRNA sequencing of the progeny of MuSCs during tail regeneration, combined with conditional regulation of key TGF-β signaling components, we identified that the level of TGF-β signaling balances the fate of MuSCs during tail regeneration.

Overall, the differentiation potential and lineage variation of MuSCs during tail and limb regeneration may represent a fundamental distinction between primary body axis and appendage regeneration in axolotls and may also extend to other vertebrate species. Exploring the intrinsic and extrinsic mechanisms governing this variation in MuSCs, and perhaps in other cell types, across these two regenerative contexts, may provide a deeper understanding to develop strategies for regenerative medicine.

## STAR⍰Methods

Detailed methods are provided in the online version of this paper and include the following:

- KEY RESOURCES TABLE
- RESOURCE AVAILABILITY

- Lead contact
- Materials availability
- Data and code availability
- EXPERIMENTAL MODEL AND STUDY PARTICIPANT DETAILS

- Axolotl husbandry
- METHOD DETAILS

- Molecular Cloning
- Transgenesis
- Drug treatment
- Whole-mount in situ hybridization
- Paraffin and cryosectioning
- Immunofluorescence
- In situ hybridization
- Single-cell transplantation
- Cell culture
- Western blotting
- Imaging
- Animal preparation and cell dissociation for Smart-seq2
- FACS and Smart-seq2 sequencing
- Sample collection and spatial transcriptomics experiment
- Smart-seq2 single-cell data process pipeline
- RNA velocity analysis and in silico genetic perturbation
- Single-cell trajectories reconstruction
- Cell differentiation status and pathway activity score prediction
- IM sub-cells identification
- Constellation plot analysis
- Spatial transcriptomics raw data processing
- Cell segmentation of spatial transcriptomics data
- Mesoderm lineage judgment
- QUANTIFICATION AND STATISTICAL ANALYSIS

## ACKNOWLEDGMENTS

We would like to acknowledge the valuable help provided by the animal facility staff Guoqing Liu and Zhenyao Wu. We thank the central equipment platform of the Guangdong Provincial People’s Hospital for its expertise and support, the China National GeneBank for providing computing resources. We also thank Dr. Prayag Murawala and Wouter Masselink for scientific discussions and valuable comments on the manuscript. This study is supported by the National Key R&D Program of China (2021YFA0805000; 2023YFA1800600; 2019YFE0106700), the National Natural Science Foundation of China (92268114; 31970782; 32070819, 32300698), the High-level Hospital Construction Project of Guangdong Provincial People’s Hospital (DFJHBF202103; KJ012021012) and BGI grant (BGIRSZ20210002).

## AUTHOR CONTRIBUTIONS

Conceptualization: J.-F.F.

Methodology: J.-F.F., Y.L., H.L., E.M.T., L.W., L.S., C.Y., J.Z., Z.Y., Y.H., X.P., N.Q., H.C., W.Z., R.Z., L.L., G.P.

Investigation: L.W., L.S., C.Y., J.Z., Z.Y., Y.H., X.P., N.Q., L.Y.

Visualization: J.-F.F., Y.L., H.L., E.M.T., L.W., L.S., C.Y.

Funding acquisition: J.-F.F., Y.L., C.Y.

Project administration: J.-F.F., Y.L., H.L., E.M.T.

Supervision: J.-F.F., Y.L., H.L., E.M.T.

Writing – original draft: J.-F.F., Y.L., H.L., L.W., L.S., C.Y.

Writing – review & editing: J.-F.F., Y.L., E.M.T., H.L., L.W., L.S., C.Y.

## DECLARATION OF INTERESTS

The authors declare no competing interests.

## Notes

### Competing Interest Statement

The authors have declared no competing interest.

